# Are there systematic biases in RNA-seq data analysis? A case study for Amphimedon queenslandica sponge as a model object

**DOI:** 10.1101/2020.02.28.969642

**Authors:** Sergey Feranchuk

## Abstract

**BACKGROUND:** The performance of a functional annotation approach for RNA-seq bioinformatics pipelines was to be compared with the method where groups of genes are generated with no relation to ontologes. Three publicly available RNA-Seq experiments for Amphimedon queenslandica sponge were used for the designed comparison. One of these experiments was referred in the publication where stages of embryo development were compared for a wide range of animal species.

**METHODS:** The expression levels were re-calculated here for three independent series of experiments. The functional annotation of differential expression levels was than conducted. This allow to compare an applicability of the two approaches, and to re-evaluate the interpretation provided in the mentioned publication.

**RESULTS:** It was confirmed by the conventional approach that Wnt and Notch pathways do operate in a development of a sponge embryo. The method of annotation which uses unbounded grouping of genes was effective in an ability to separate development stages of sponge embryo. In addition, the published results were by a suggestion distorted by an artifact, caused by a positive feedback in the stage of data processing.

## Introduction

There are ambiguities in a choice of pipeline for bioinformatics processing of RNA-seq experiments. Data to be processed, besides of a setup of the experiment, do always depend on multiple stresses and causalities.

Functional annotation of differentially expressed genes is one of the issues where a presence of this choice is underestimated. A lot of RNA-seq projects were successfully completed due to hints from functional annotation. But this is just another reason to put an attention to an extent of its applicability, and to look around to seek any alternative approaches.

A case study can demonstrate limitations of that approach more definitely. A baseline tools of functional annotation are anyway successful, and usefulness of it can be argued only when particular precedents are examined.

A sponge Amphimedon queenslandica was selected as a model for the case study by an occasion. This object has some specific features, as well as many other biological objects do.

A sponge is distant from most of carefully described animal species. An interpretation of its genomic data has additional obstacles, unusual and unexpected in conventional pipelines.

Anyway, three independent series of RNA-seq experiments are available in NCBI for A.queenslandica, which makes it possible to conduct a correct and complete comparative research. The setup in all of these experiments was declared to be focused on a development of sponge embryo.

At the moment, only one project for the one series of experiments has references to a completed publication. The results of this project were reported in methodological study [2] and in the study [3], where embryonic development of animal species was compared.

In this situation, in order to discuss an applicability of functional annotation, the major statement of the referred study should also be commented. This statement was in a strong enough suggestion, that a certain point of divergence did exist in a development of all phyla of animal kingdom.

The major warning which can by expectation be caused by an over-estimated power of functional annotation is a presence of a “positive feedback”, gradually accumulated declinations in opinions obtained this way. By an occasion, the over-strengthened major statement of the article [3] can be relaxed, if to assume a presence of a “positive feedback” in an over-complicated methodology of data processing, which was developed to support the declared results.

### The approach

The confidence of a statistical hypothesis at present is the only neutral criteria which can verify the interpretation of experiments on differential expression of genes. Sometimes this allows to select some particular genes which are expressed differently in the compared groups of samples.

In other cases, an acceptable value of statistical significance can be obtained when genes in some group are expressed differently. An assumption is implied here that these genes are definitely connected to the group by some certain criteria.

The matter is that “gene” itself means usually a complex and even not completely certain biological entity, and a lot of criteria can be used to join some genes into a group. Some of criteria and some of groups appear to be more efficient and appropriate for heterogeneous experiments. But a “positive feedback” can be assumed on this stage, the proved importance of a term in the previous investigations will motivate the choice in favor to that term instead of more appropriate but less known groupings.

Another analysis of differential expression is presented here for a comparison; it was based on neutral unbounded groups of genes. The groups were generated as sub-communities in a network of gene interactions, and a random component was introduced in the algorithm of community detection.

The purpose of the presented case study was to demonstrate the usefulness of that another approach and to compare an extent of applicability of the two approaches in an analysis of differentially expressed genes. The examined comparison allows to show, in which way the assumed positive feedback can appear.

### Previous works

The embryo of sponge pass seven stages before transforming to a free-swimming larvae. A stage of embryo development can be detected visually.

The development of sponge embryo attracts an attention because it can expand the understanding about the generic pathways which operate in a development of multicellular animal embryos. It was supposed and observed that the signaling pathways do in some way operate in sponge, which determine a formation of body in embryo of most animals and are associated with definite families of proteins, Wnt signaling proteins and Notch receptors [1].

A more strong result was reported in [3] which was obtained relying on a comparison of the expressed transcripts for 35 species from different animal phyla. There was shown, that a phase of transition does exist in a development of any embryo, and all the genes in a genome can be separated into the three groups – one group operates mostly in an early development of embryo, another group in later stages, including adults, and a third group is active mostly in a period of the transition phase.

The presence of that transition phase in an embryo development of all animals was demonstrated there using comparison of gene expression profiles at different stages. It was suggested also that a presence of this transition phase is accordant on a time scale with the divergence of animal species in early evolution of life.

As an experimental proof of this hypothesis, total RNA was extracted and sequenced, for separate embryo cells in different phases of development. In this setup of the experiment, there is a need to assign and verify an exact stage of development for each of the sequenced samples.

An approach was proposed in [2] which do implement this ordering of samples, by resolving a problem of combinatorial optimization. The selection of a criteria for optimization was based on an idea that more genes are actively expressed in later phases of embryo development.

The results of that comparative analysis of transcriptomes were shown to be consistent with previously known observations concerning embryo development. In particular, genes associated with Wnt and Notch signaling pathways appear to be abundant in the measured transcriptomes, with a significance even below 0.001, the estimated significance for A.queenslandica was around 0.1.

## Methods

The surveys which were used in the comparative study are listed in table 1. Illumina HiSeq 2000 sequencing with single-end reads was used in all three surveys.

**Table 1.**
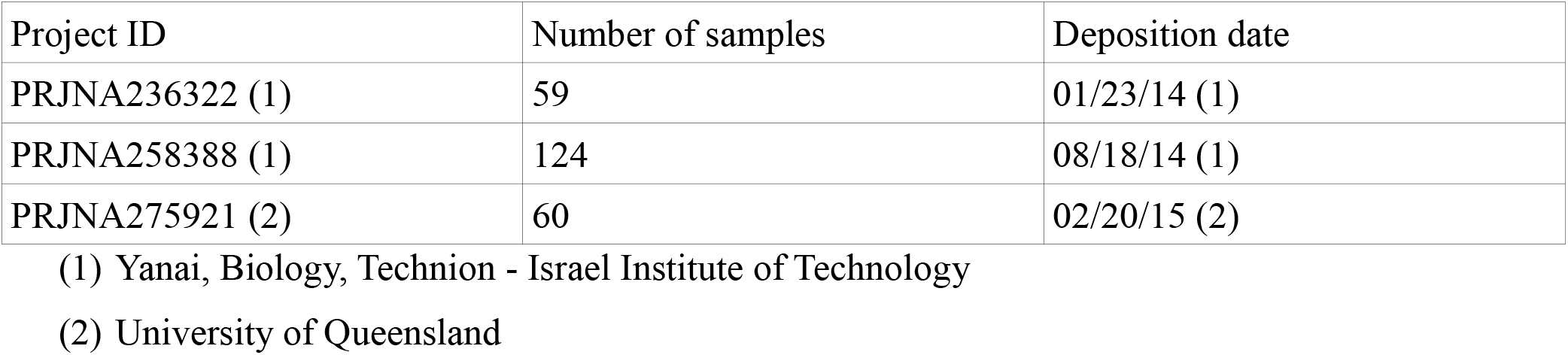

| Project ID | Number of samples | Deposition date |
| --- | --- | --- |
| PRJNA236322 (1) | 59 | 01/23/14 (1) |
| PRJNA258388 (1) | 124 | 08/18/14 (1) |
| PRJNA275921 (2) | 60 | 02/20/15 (2) |

(1) Yanai, Biology, Technion - Israel Institute of Technology
(2) University of Queensland

Coding sequences of genes were used to estimate levels of gene expression from the transcriptome sequencing. To do this, raw archives of reads were downloaded from NCBI and pre-processed using Trimmomatic software. After that, the reads were aligned to Aqu2 version of A.queenslandica genome with bowtie2 and a median coverage of each gene sequence was transformed to a level of expression.

In order to generate an array of loosely bounded groups of genes, a network of gene interactions was reconstructed based on the estimated levels of gene expression. Edges in the network were assigned if a value of correlation between genes was above 0.7. Kendall rank correlation was applied there, for all the samples from the three surveys. To generate 1000 groups of genes, a sub-network of a randomly selected genes was separated each time, and a combination of community detection algorithms from “igraph” software library was applied to this sub-network. In the result, the sizes of obtained groups were around 16, and strictly within interval 16±1.

The samples for stage 1 (cleavage), stage 2 (brown) and stage 4(spot), from the three surveys, were used for a comparative presentation. The differentially expressed genes, for these three stages, were selected, with a p-value of significance below 0.001. The significance on this step was estimated by Anova criterion, implemented in scipy python library. For any group, some genes in it were observed to be differentially expressed, and a number of these genes could show the applicability of this group for a differentiation of samples. The number of genes was transformed to a level of significance by the exact Fisher test, as a measure of enrichment.

This criteria of enrichment was used to demonstrate a fitness of a group to a particular sample, and to select groups which most efficiently represent the overall separation of samples. Pearson measure of correlation, PCA decomposition implemented in sklearn library and “permanova” algorithm from skbio library were used to re-evaluate and demonstrate the efficiency of the separation, for the three selected groups. The estimated p-values were of order above *e*^-4^ for the annotated groups of genes and of order around *e*^-6^ for the selected unbounded groups.

Interproscan pipeline of functional annotation was used to select genes related to Wnt and Notch pathways, from the A.queenslandica genes. Charts in the figures were prepared with Matplotlib python library and Inkscape program was than applied to improve the visualizations. A script on python was developed to demonstrate an appearance of an artifact in a simulation of a RNA-seq pipeline. Source codes of the scripts and intermediate data were deposited to Zenodo and are available at doi://10.5281/zenodo.3690221

## Results

The lineage of sponges is distant from other animal phyla, and genes of sponges are distant by a structure and a function from their homologs in other model species. Possibility to annotate sponge genes is limited; only five genes related to Wnt pathway and six genes related to Notch pathway were found in the results of the functional annotation. These genes were used to present an enrichment of the corresponding groups which is shown in fig 1.

**Fig. 1.**
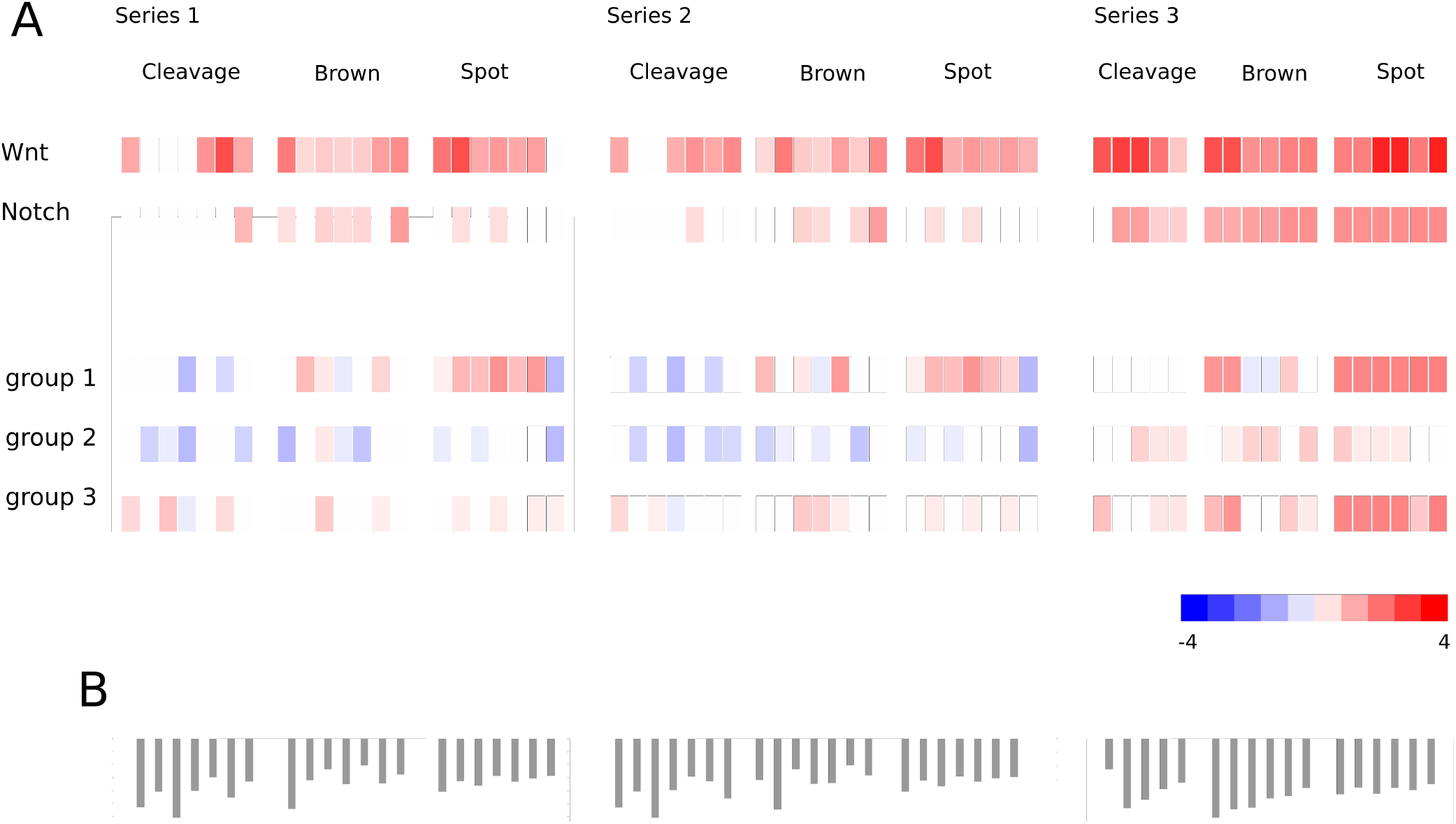
Heatmap for an enrichment of gene groups in the series of samples. A value of enrichment is estimated as a p-value of Fisher exact test applied to a number of differentially expressed genes in a sample. The color map corresponds to natural logarithm of the p-value. Bars on the part B show a number of genes with an expression level above average.

These genes are indeed abundantly expressed in most of the samples, and levels of their expression vary a lot from sample to sample. The evidences of common origins in an embryo development of diverged species are therefore observed and it anyway can not be doubted.

A collected ontology which concern these signal pathways appears to be confirmed. And the usability of conventional approaches of functional annotation is demonstrated for this case.

Another interest in the setup of the analysis is to detect definite changes in gene expression for different stages of development. For overall gene expression profiles, a straightforward PCA decomposition of correlation matrix doesn’t allow to separate the development stages (Fig. 2A). The distribution of genes in the two annotated groups of genes is also no enough correlated with a stage of development.

**Fig. 2.**
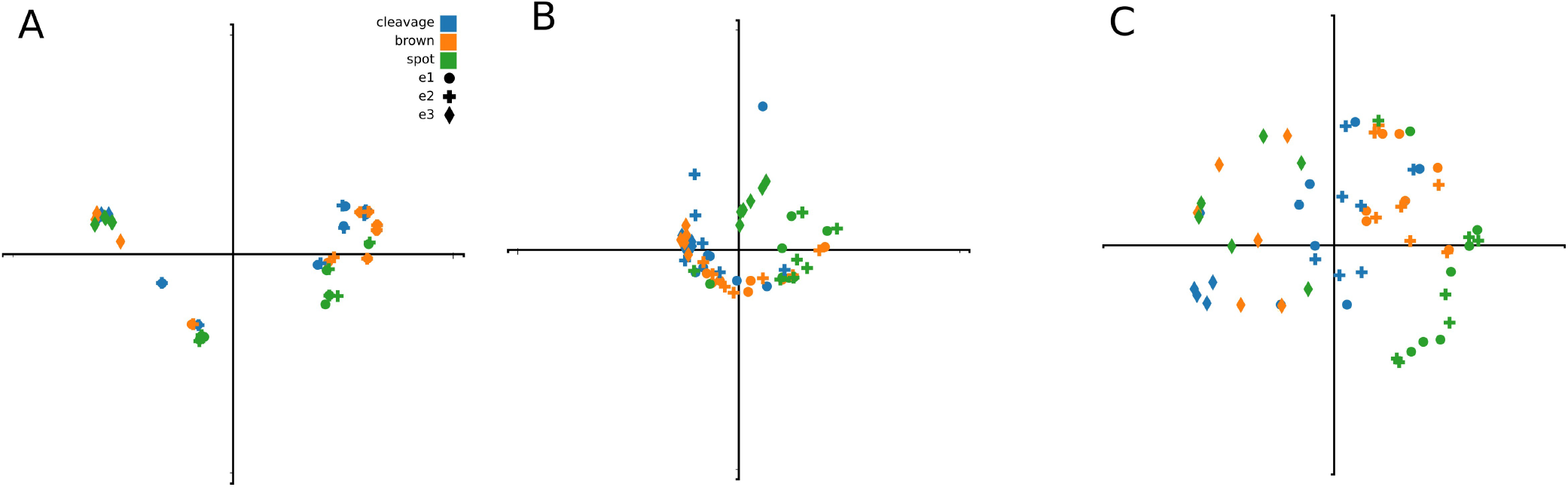
PCA decomposition of correlation matrix, for overall gene profiles (A) and for genes in groups 1 and 2 (B,C).

Several groups of genes, selected from 1000, separate the stages with a better significance, and this allows to reduce the distinguishing of development stages to an analysis of several dozens of genes (Fig. 2B, 2C). Although these groups were chosen as appropriate only in the specific setup of the separation, the proposed approach does put forward the investigation of sponge embryo development; a possibility is provided to focus a subject of that research on genes in the selected groups.

A relative number of genes which were expressed in a sample, shown in fig. 1B, is not increased in later stages of development. This is in accordance with the results from [3] where a sufficient number of genes were included in a group which was specific to early stages of an embryo development.

And in that way a presence of the artifact can be supposed in the methods of data processing reported in [2], so that the major statement of the article can be doubted. The feature of ‘reversed bottleneck’ which was stably observed in the provided comparisons of transcriptomes can be explained either as a phenomenon in the evolution of species or due to a presence of a positive feedback in the data analysis.

An appearance of similar artifact on an artificial example is demonstrated in fig 3; the reordering applied to the samples was analogous by a setup to one used in [2]. A spot of samples with a negative correlation is stably enough condensed in a right corner of the heatmaps after reordering.

**Fig. 3.**
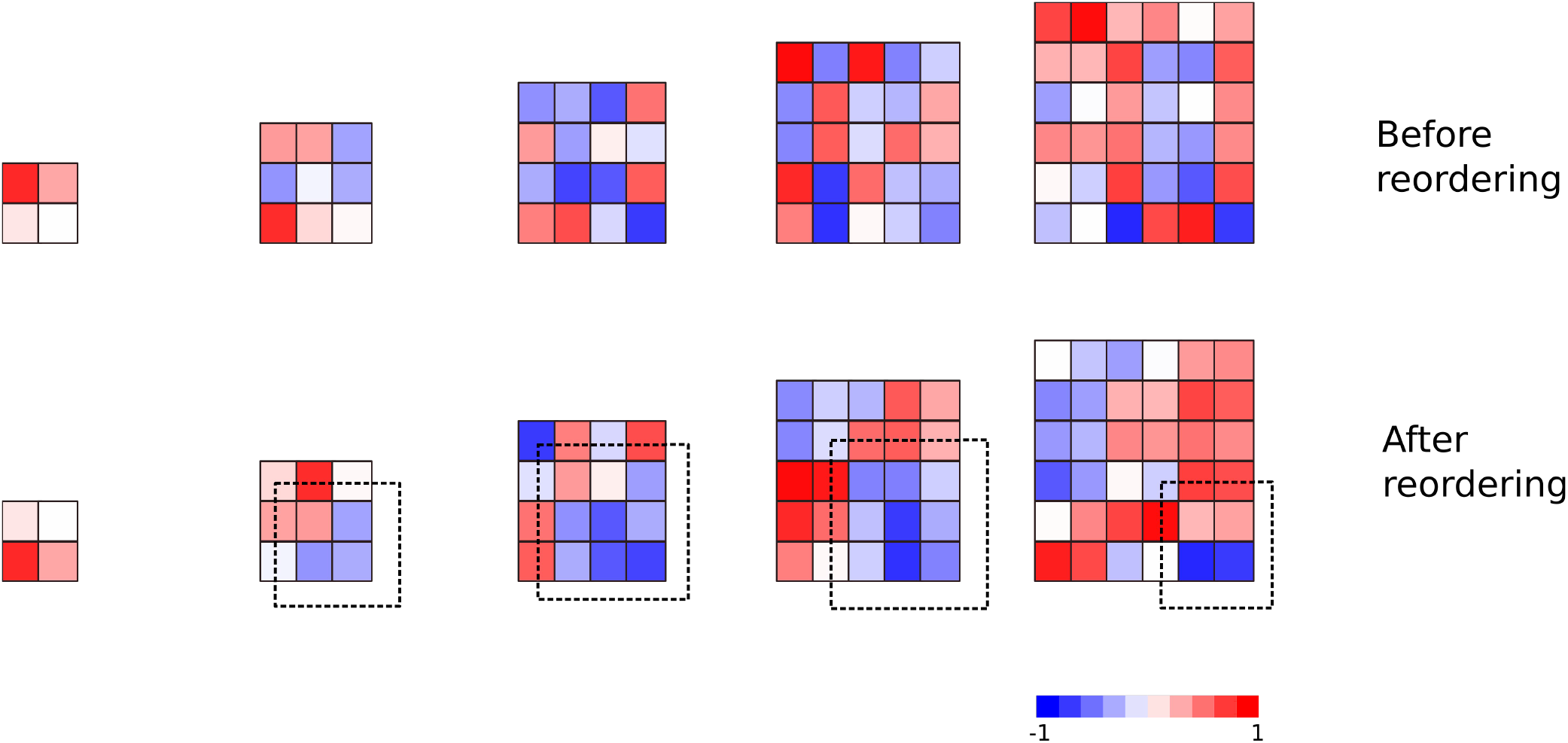
An artificial example which illustrates the appearance of an artifact. Points on a heatmap show a value of correlation between expression levels in the simulated samples. All the samples were combined from 6 “genes”, levels of their “expression” were randomly shuffled in each sample. In a bottom part, the samples were reordered following the level of 1-st “gene” (vertical series) and 2-nd “gene” (horizontal series).

A way in which a pairs of transcripts were compared can also introduce a distortion to the methods; it was observed in intermediate tests that when one transcript from a first series is paired with many others in a second series, changes in the resulted image are clearly seen. Anyway, the presence of positive feedback in the methods is almost certain, and this can explain the repeated appearance of the ‘bottleneck’ in correlation heatmaps, no exceptions from this rule were reported.

## Discussion

The provided comparison of grouping methods confirms the usability of the conventional approach in which grouping of genes follows the collected ontology. This direction is productive and its advances are expected to be productive too.

And another approaches are expected to be effective to resolve particular applied problems, the usability of one of such approaches was demonstrated above. Quit different solutions can be invented for some specific cases, to analyze RNA-seq data.

There are the reasons to suggest a presence of the artifact in the method called “Blind”, which was used to re-order the samples in the study [2], and to doubt the major statement of that article. This statement sounds as too ‘sensational’ and unexpected, and it was doubted by other scientists [4].

This artifact is caused by a presence of “positive feedback” in the method. The model which was assumed for the validation of the method was not applicable to the experimental data.

Is a usability of ontology grouping over-estimated? A gene enrichment analysis is often supported by a solid credibility of p-values.

Probability distributions with infinite dispersion are most often suitable to describe living systems. The methods where credibility is estimated in p-value are not applicable to these distributions.

It can cause a positive feedback, an appearance of an artifact. Errors of this kind can be associated with infinite loops in programming, or incidents in behavior when a patient repeat the same actions when quit another strategy should be chosen.

## Acknowledgments

The author is grateful to former colleagues in Limnological institute in Irkutsk, who motivated his interest to the subject.

